# Sensory uncertainty punctuates motor learning independently of movement error when both feedforward and feedback control processes are engaged

**DOI:** 10.1101/2022.09.01.506147

**Authors:** Christopher L. Hewitson, David M. Kaplan, Matthew J. Crossley

## Abstract

Integrating sensory information during movement and adapting motor plans over successive movements are both essential for accurate, flexible motor behavior. When an ongoing movement is off target, feedback control mechanisms update the descending motor commands to counter the sensed error. Over longer timescales, errors induce adaptation in feedforward planning so that future movements become more accurate and require less online adjustment from feedback control processes. Both the degree to which sensory feedback is integrated into an ongoing movement and the degree to which movement errors drive adaptive changes in feedforward motor plans have been shown to scale inversely with sensory uncertainty. However, since they have only been studied in isolation of each other, little is know about how they respond to sensory uncertainty in real-world movement contexts where they co-occur. Here, we show that sensory uncertainty impacts feedforward adaptation of reaching movements differently when feedback integration is present versus when it is absent. In particular, participants gradually adjust their movements from trial-to-trial in a manner that is well characterised by a slow and consistent envelope of error reduction. Riding on top of this slow envelope, participants display large and abrupt changes in their initial movement vectors that clearly correlate with the degree of sensory uncertainty present on the previous trial. However, these abrupt changes are insensitive to the magnitude and direction of the sensed movement error. These results prompt important questions for current models of sensorimotor learning under uncertainty and open up exciting new avenues for future exploration.

**Author Summary:** A large body of literature shows that sensory uncertainty inversely scales the degree of error-driven corrections made to motor plans from one trial to the next. However, by limiting sensory feedback to the endpoint of movements, these studies prevent corrections from taking place during the movement. Here, we show that when such corrections are promoted, sensory uncertainty punctuates between-trial movement corrections with abrupt changes that closely track the degree of sensory uncertainty but are insensitive to the magnitude and direction of movement error. This result marks a significant departure from existing findings and opens up new paths for future exploration.

## Introduction

During episodes of sensorimotor control, sensory information about the current state of the body and environment are used to generate motor commands to achieve a desired goal. In an ideal world, this process would be implemented perfectly and result in error-free motor behavior. In the real world, however, every stage of the sensorimotor control process is contaminated by noise [1] and uncertainty [2]. Despite this, humans achieve remarkably accurate and appropriate motor behavior by harnessing two complementary processes for error correction. First, sensory feedback is rapidly integrated to adjust ongoing movements and compensate for sensed deviations from the planned movement [3, 4, 5, 6, 7]. Second, over successive movements, feedforward motor plans, which map behavioral goals to the motor commands needed to accomplish those goals, are adapted in response to movement errors [6, 8, 9, 10, 11, 12, 13]. Throughout this paper, we refer to the former as *feedback integration* and the latter as *feedforward adaptation*.

An important question that has attracted considerable attention recently concerns how these error correction processes are influenced by sensory uncertainty. To date, this issue has primarily been investigated using experimental paradigms that isolate feedback integration from feedforward adaptation, or vice versa. For example, in their well-known study, Körding and Wolpert [14] showed that visual feedback provided briefly at the midpoint of the reach drives movement corrections that are inversely proportional to the level of uncertainty in the sensory feedback. In other words, when uncertainty is high, sensory feedback information is integrated less to correct the ongoing reach (and reliance on prior knowledge increases) compared to when visual uncertainty is low. Several follow-up studies have made similar observations [15, 16]. Studies investigating the feedforward adaptation component have similarly shown that adaptation rates inversely scale with sensory uncertainty such that increasing sensory uncertainty leads to smaller updates to the feedforward plan (slower adaptation) and vice versa [15, 17, 18, 19, 20, 21, 22].

Importantly, because the majority of studies investigate these processes in isolation, little is known about how sensory uncertainty influences feedback integration and feedforward adaptation when they co-occur – as they do in most natural movement contexts. For example, if highly uncertain sensory feedback leads to relatively small online corrections during a movement, does it also drive similar adaptive changes in feedforward motor plans? To our knowledge, no existing studies address this key question.

Here, we examine how sensory uncertainty influences feedforward adaptation and feedback integration when they co-occur. Our results indicate that (1) the presence of feedforward adaptation has little to no effect on how sensory uncertainty influences feedback integration, but (2) in the presence of feedback integration, sensory uncertainty appears to punctuate a slow and steady envelope of error reduction with large and abrupt changes to initial movement vectors that are insensitive to the magnitude and direction of the sensed movement error. This latter finding represents a significant departure from the existing literature which consistently reports that sensory uncertainty inversely scales an error-dependent response.

## Results

The overarching aim of our experiments is to identify how feedforward adaptation and feedback integration interact in the response to different levels of sensory uncertainty. Participants made planar reaching movements on a low friction surface (see Fig. 1) with feedback about their hand position provided only at movement onset, midpoint, and / or endpoint. Only feedback at midpoint can influence feedback integration, while feedforward adaptation may be influenced both by feedback at midpoint and endpoint.

**Figure 1.**
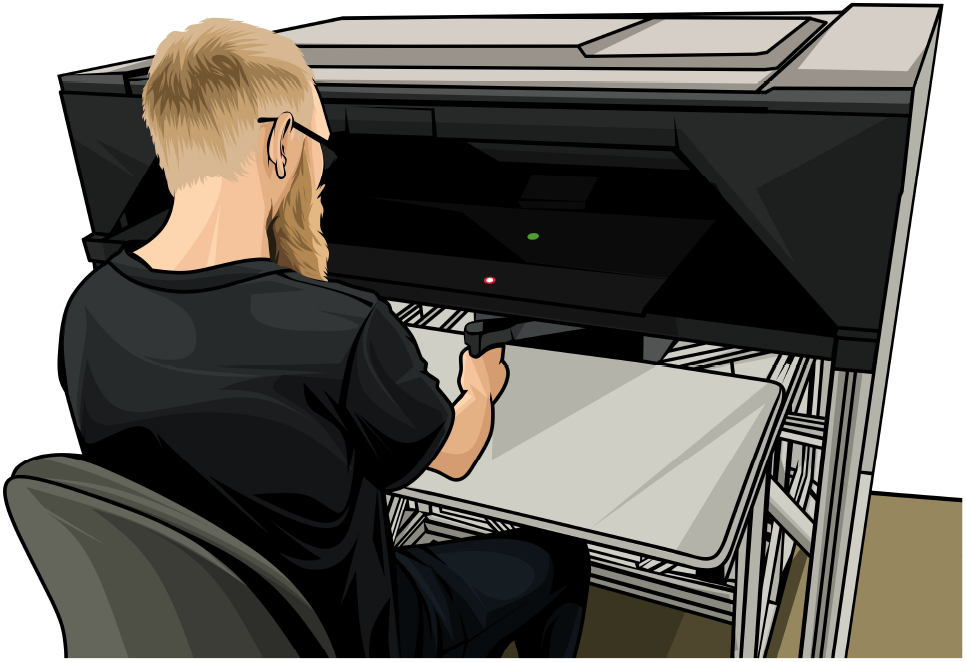
Kinarm Endpoint Robot. Green start target, red end target, and white cursor feedback are represented.

### Experiment 1

The primary contribution of Experiment 1 is to identify how feedforward adaptation responds to sensory uncertainty when feedback integration is also engaged. We provided uncertain sensory feedback at movement midpoint (see Fig. 2A), and omitted endpoint feedback entirely. We focused on uncertainty at midpoint because (1) feedback integration can occur only if feedback is provided at some point before movement endpoint, and (2) it is likely that midpoint feedback alone is sufficient to drive feedforward adaptation [14], so (3) including only midpoint feedback is the simplest possible design in which feedback integration and feedforward adaptation can co-occur.

**Figure 2.**
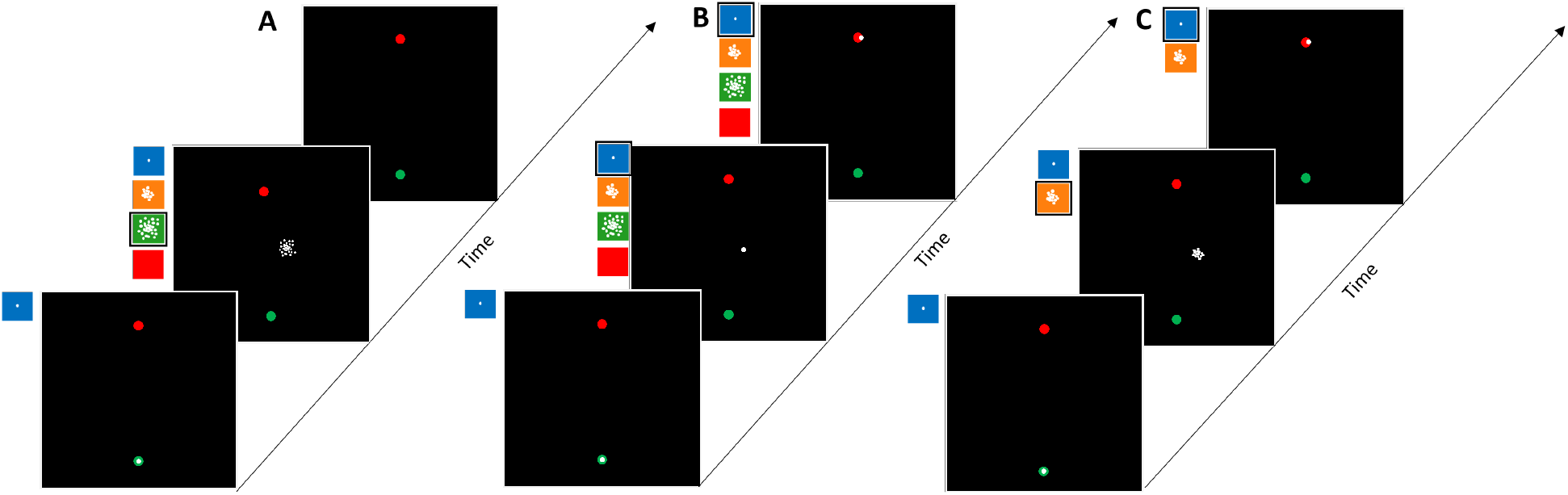
Experimental protocols: Example adaptation phase trial conditions. (A) Experiment 1: Midpoint only feedback (large uncertainty). (B) Experiment 2: Matched midpoint and endpoint feedback (low uncertainty). (C) Experiment 3: Unmatched midpoint (moderate uncertainty) and endpoint (low uncertainty) feedback. Bottom, middle and top slides represent start, middle and end of reach respectively. Colored panels represent the possible uncertainty conditions (blue: σ_0_, orange: σ_M_, green: σ_H_, red: σ_∞_). The example condition applied is outlined in black. In all experiments, a no-feedback washout phase followed the adaptation phase.

Figure 3 shows group-averaged initial movement vectors per trial color coded in panel A such that the color of the dot at trial *t* indicates the sensory uncertainty experienced at midpoint on trial *t* – 1, and in panel B such that the color of the dot at trial *t* indicates the perturbation applied on trial *t* – 1. Recall that the trial sequence of perturbations and sensory uncertainty levels was matched across participants, which is a fundamental design feature that makes this plot informative. There are a few important features to take from Figure 3.

**Figure 3.**
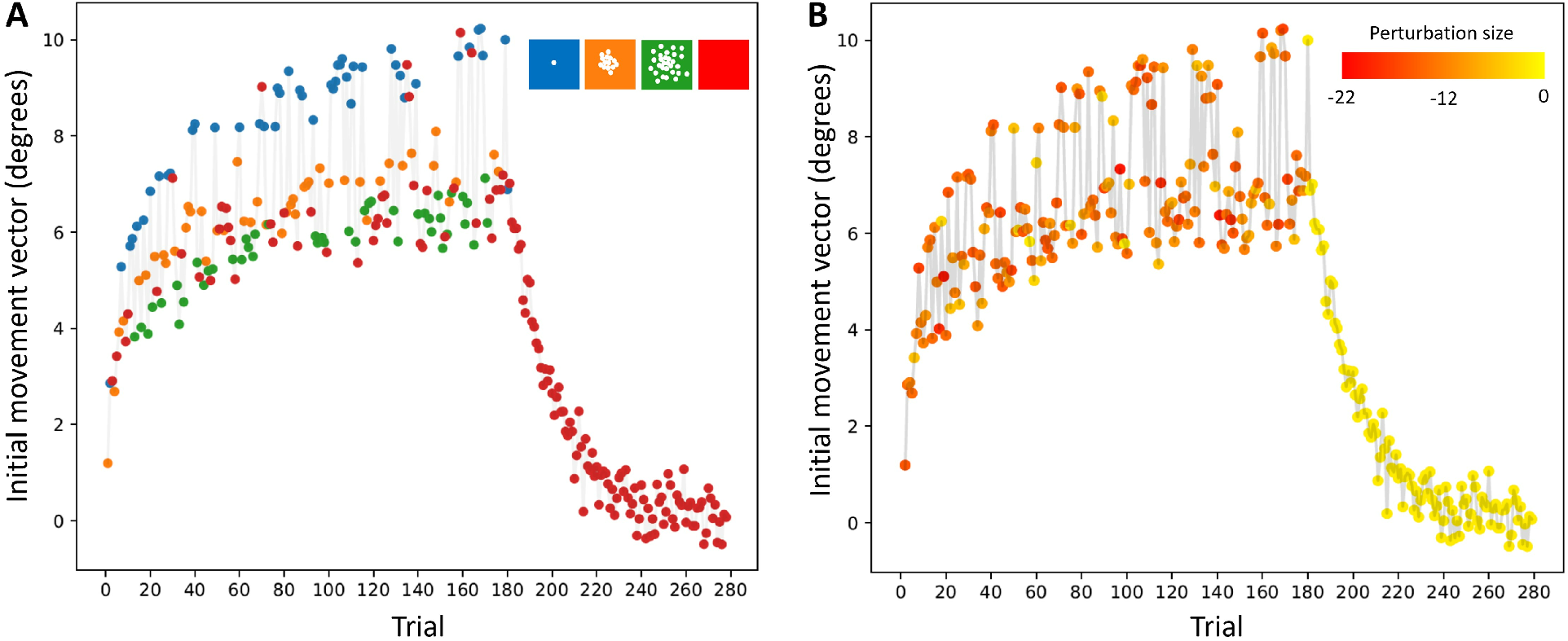
Mean initial movement vector across participants per trial in Experiment 1. (A) The color of the dot on trial t represents the level of sensory uncertainty applied at midpoint on the previous trial t – 1. (B) The color of the dot on trial t represents the perturbation applied on the previous trial t – 1.

First, the change in initial movement vectors across trials during the adaptation phase is in a direction that – on average – tends to reduce error. Second, note that initial movement vectors decay smoothly back to baseline during the washout phase. Together, these observations are consistent with the idea that changes in initial movement vector across trials are driven – on average – by an incremental adaptive process.

Second, the trial-by-trial variation in initial movement vector is quite large, but this variation is not due to noise. Panel B suggests that these large trial-wise fluctuations in reaching behavior are not well correlated with the sequence of perturbations – and are therefore not well correlated with the magnitude of the movement errors – experienced across the experiment. Rather, as shown in panel A, there is a clear stratification across sensory uncertainty experienced at midpoint. Thus, sensory uncertainty at midpoint appears to have a dramatic and systematic effect on the evolution of initial movement vectors across trials.

In particular, note that any change across trials from a lower uncertainty level to a higher uncertainty level (e.g., *σ_L_* → *σ_M_*, *σ_M_* → *σ_H_*) almost always results in a change in movement vector that is anti-adaptive. To understand this, recall that the perturbation experienced on trial *t* is on average 12°, and therefore the error experienced on every trial should drive the subsequent trial’s initial movement vector more in the positive direction. Instead, we see on these trials, initial movement vectors are driven back down in a direction that leads to greater error on the subsequent trial.

Considered the other way around, any change across trials from a higher uncertainty to a lower uncertainty level (e.g., *σ_M_* → *σ_L_, σ_H_* → *σ_M_*) almost always results in a change in movement vector that is adaptive (i.e., in an error-reducing direction) but of a much greater magnitude than would be seen if the uncertainty level did not change at all.

Figure 4 shows a comparison of one-state and two-state model variants. Panels A-C show initial movement vectors overlaid with one-state model predictions. All one-state models fit poorly to the behavioural data. The reason these one-state models perform so poorly on our data is that they have no means by which to capture the trial-by-trial uncertainty-dependent fluctuations that go against the error-reduction gradient. The best they can do is to retain essentially nothing from trial-to-trial as this allows any accrued adaptation on trial *t* to be forgotten on subsequent trial *t* + 1, which can in some cases can allow for changes in hand angle that go against the error-reduction gradient. In line with this, the best-fitting retention parameter for the one-state models is zero for every participant. This is also why the model predictions during washout fall so suddenly to baseline, which further hurts the quality of single-state model fits. From these results, it is clear that single-state models cannot provide a good fit to our behavioural data.

**Figure 4.**
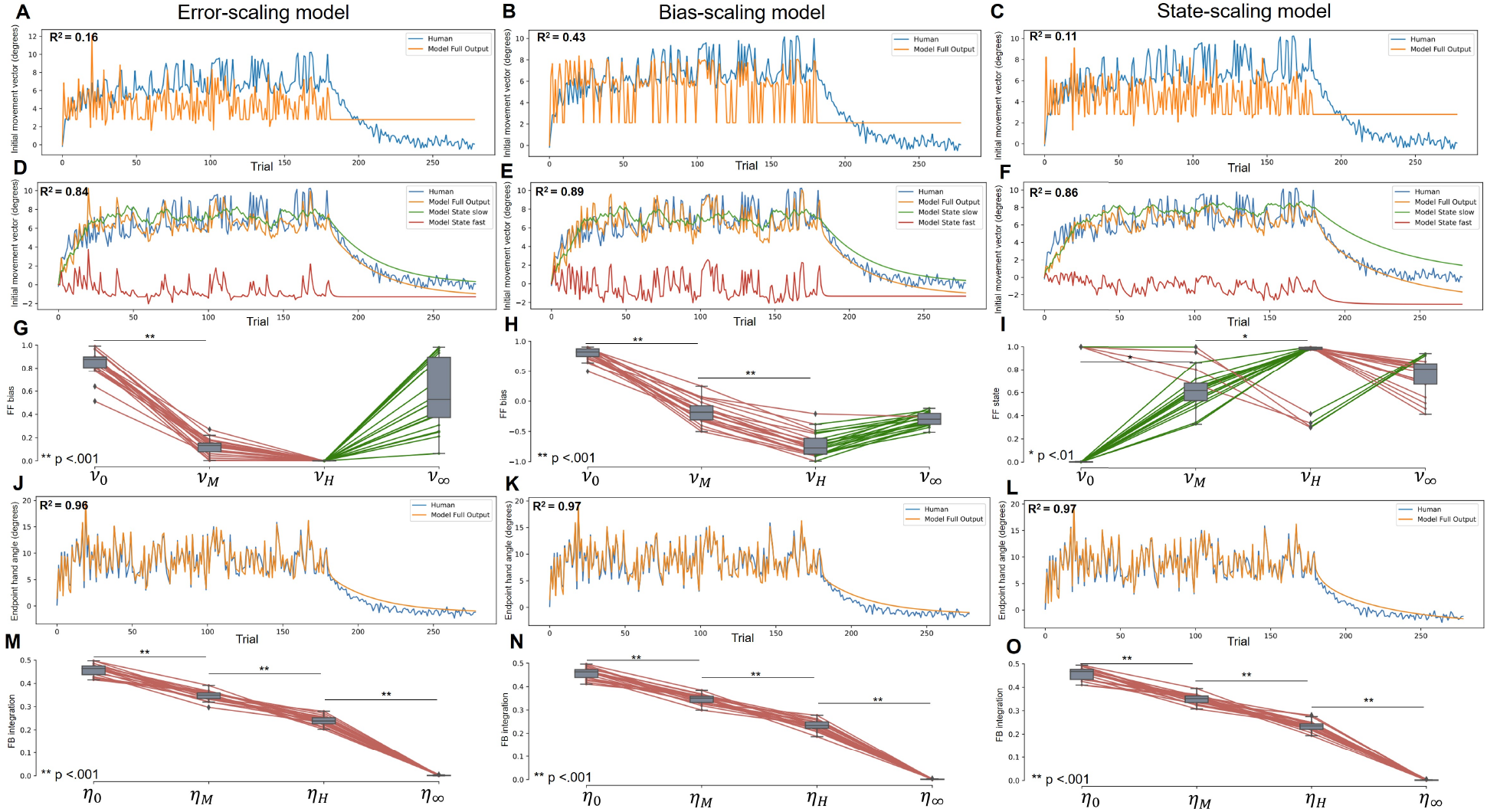
Panels A-C show initial movement vectors averaged across participants in Experiment 1 overlaid with that predicted by the (A) one-state error-scaling model, (B) one-state bias-scaling model, and (C) one-state state-scaling model. Here, human performance averaged across participants is shown in blue and model predictions are shown in orange. Panels D-F show initial movement vectors averaged across participants overlaid with that predicted by the (D) two-state error-scaling model, (E) two-state bias-scaling model, and (F) two-state state-scaling model. Here green and red traces show averaged slow and fast model components separately, which combine to produce the full model predictions in orange. Panels G-I show the distribution of best fitting **ν** parameters for the (G) two-state error-scaling, (H) two-state bias scaling, and (I) two-state state-scaling model. Here, individual differences between parameters are represented by a single colored lines that are green when parameter values increase and red when they decrease. Panels J-L show endpoint hand angles averaged across participants overlaid with that predicted by the (J) two-state error-scaling model, (K) two-state bias-scaling model, and (L) two-state state-scaling model. Here, human performance averaged across participants is shown in blue and model predictions are shown in orange. Panels M-O show the distribution of best fitting **η** parameters for the (M) two-state error-scaling, (N) two-state bias-scaling, (O) two-state state-scaling model.

Panels D-O show a comparison of two-state model variants. Here, the behavioral data overlaid with model predictions and best-fitting parameters. All two-state models capture the data in a qualitatively similar way. In particular, the slow system state variable (green line in panels D-F) carries the overall adaptation envelope across trials, and the fast system hovers around zero with large fluctuations across trials. From visual inspection, the bias-scaling model does a better job tracking the large uncertainty-dependent fluctuations across trials than either the error-scaling model or the state-scaling model. In line with this, the error-scaling model (*R*^2^ = 0.84) accounts for less of the variance in the behavioural data than either the state-scaling model (*R*^2^ = 0.86) or the bias-scaling model (*R*^2^ = 0.89).

Model comparison via Δ*BIC* reveals that the single-state models perform significantly worst than their two-state counterparts: single-state vs two-state error-scaling Δ*BIC* = −224.8 ± 3.4, *t*(38) = −67.1, *p* > .001, *d* = − 21.2); single-state vs two-state bias-scaling Δ*BIC* = −189.9 ± 6.3, *t*(38) = −41.4, *p* > .001, *d* = −9.5); and single-state vs two-state state-scaling Δ*BIC* = − 237.1 ± 3.5, *t*(38) = −68.1, *p* > .001, *d* = −21.52).

Comparing two-state models via Δ*BIC* provides very strong evidence that the bias-scaling model best accounts for the data in all 20 participants (bias-scaling vs error-scaling Δ*BIC* = −23.7 ± 4.1, *t*(38) = 5.7, *p* > .001, *d* = 1.8; bias-scaling vs state-scaling Δ*BIC* = −19.1 ± 4.2, *t*(38) = 4.5, *p* > .001, *d* = 1.4).

A one-way ANOVA with best-fitting *ν* values from the bias-scaling model as the factor indicates that sensory uncertainty significantly influenced the bias term applied to the feedforward update (*F* = 634.6, *p* < .001, *ω*^2^ = 0.93). Posthoc t-tests showed that sensory uncertainty inversely scaled the feedforward adaptive update (*ν*_0_-*ν_M_* : *t*(38) = 20.4, *p* < .001, *d* = 5.1; *ν_M_-ν_H_* : *t*(38) = 20.1, *p* < .001, *d* = 4.5). Interestingly, while the expected order of sensory uncertainty scaling is observed across the feedback conditions (i.e *ν*_0_ > *ν_M_* > *ν_H_*), the no-feedback condition (*ν*_∞_) parameter is not significantly different to the high-uncertainty condition (*ν_H_-ν_∞_* : *t*(38) = −10.4, *p* = .99, *d* = −2.3), and not significantly lower as expected.

In summary, when driven by midpoint feedback – and thus when co-occurring with feedback integration – feedforward adaptation is not well captured by the error-scaling model. This challenges the dominant view that sensory uncertainty inversely scales an error-dependent feedforward adaptive update term.

The influence of sensory uncertainty on feedback integration, on the other hand, appears to be well captured by the traditional error-dependent, inverse scaling by sensory uncertainty model [14]. In particular, the feedback integration aspect of the model lead to an *R*^2^ = 0.97 in model endpoint compared to human endpoint data. A one-way ANOVA with best-fitting *η* values from the best fitting bias-scaling model as the factor confirms that there are differences in how much feedback was integrated across different uncertainty levels (*F* = 1892.5, *p* < .001, *ω*^2^ = 0.98). Posthoc t-tests indicate that indeed *η*_0_ > *η_M_* > *η_H_* > *η_∞_* (*η*_0_ – *η_M_* : *t*(38) = 16.1, *p* < .001, *d* = 3.6, *η_M_* – *η_H_* : *t*(38) = 15.6, *p* < .001, *d* = 3.5, *η_H_* – *η_∞_* : *t*(38) = 47.6, *p* < .001, *d* = 10.7).

### Experiment 2

Experiment 1 revealed that midpoint feedback is sufficient to drive feedforward adaptation, but that the uncertainty of this feedback does not inversely scale the magnitude of the error-driven feedforward update as it does in an endpoint feedback-only paradigm [17, 18, 19, 20, 21, 22]. Rather, it acts independently of the experienced error to induce large and abrupt changes in the adapted state that can be – in the case of a transition from a high uncertainty trial – in an anti-adaptive direction.

It is therefore possible that the presence of feedback integration fundamentally alters how the feedforward adaptation process is affected by sensory uncertainty. However, another possibility is that feedforward adaptation is influenced by sensory uncertainty differently depending on the temporal proximity of the sensory feedback signal to movement offset, regardless of whether or not feedback integration occurred at the midpoint of the movement [23, 24, 25]. Experiment 2 tests this possibility by providing midpoint and endpoint feedback matched in their level of uncertainty (see Fig. 2B).

Figure 5 shows group-averaged initial movement vectors per trial color coded such that the color of the point at trial *t* indicates the sensory uncertainty experienced at midpoint on trial *t* – 1. The same basic pattern of results observed in Experiment 1 (Figure 3) is present here as well. Specifically, the trial-by-trial variation in initial movement vector is quite large, but cannot be attributed to noise because there is a clear stratification in the how a particular uncertainty level influences the subsequent trial. Furthermore, we see again that in the transition from lower uncertainty trials to higher uncertainty trials (e.g., *σ_L_* → *σ_M_*, *σ_M_* → *σ_H_*), the change in movement vector is anti-adaptive.

**Figure 5.**
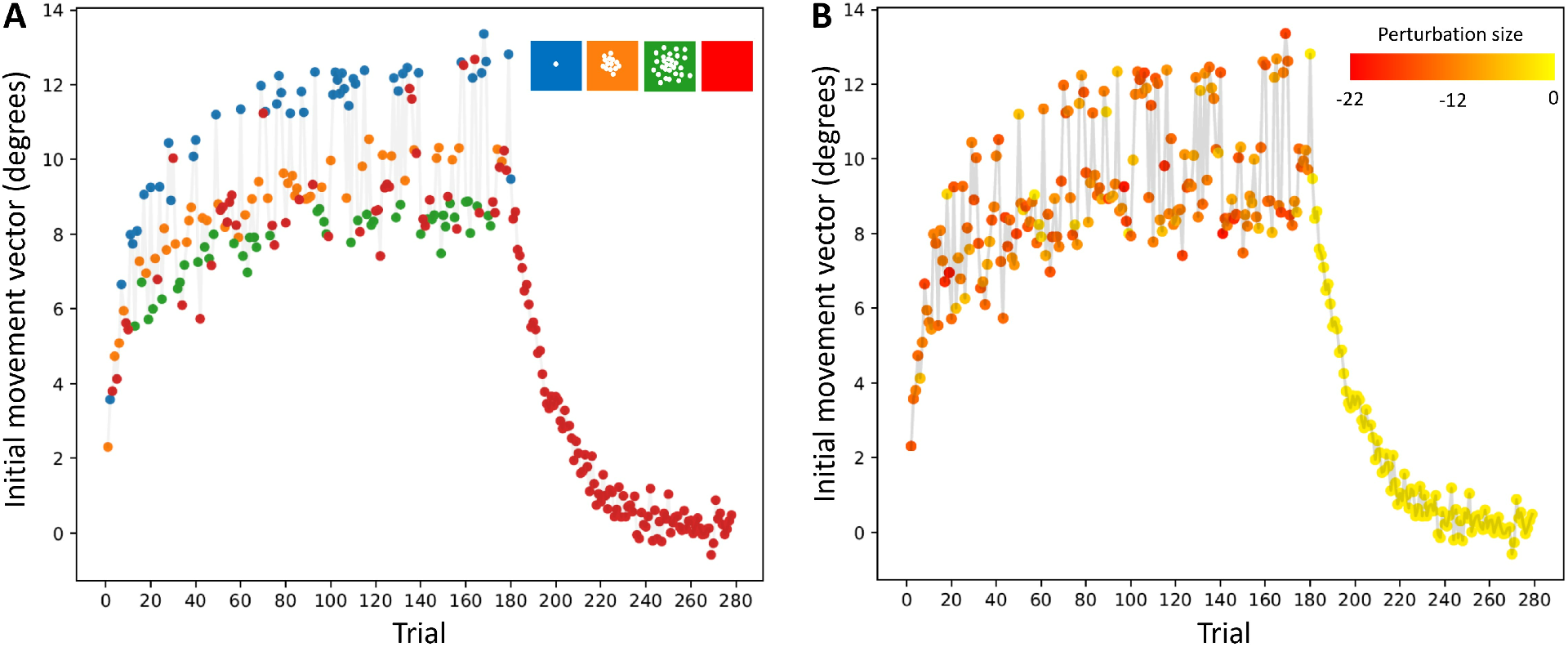
Mean initial movement vector across participants per trial in Experiment 2. (A) The color of the marker on trial t represents the level of sensory uncertainty applied at midpoint on the previous trial t – 1. (B) The color of the marker on trial t represents the perturbation applied on the previous trial t – 1.

Figure 6 shows a comparison of one-state and two-state model variants. Here, the behavioral data is overlaid with model predictions and best-fitting feedforward and feedback parameters. Panels A-C show initial movement vectors overlaid with one-state model predictions. All one-state models fit poorly to the behavioural data as seen in experiment 1. Panels D-O shows a comparison of two-state model variants. Here, the behavioral data overlaid with model predictions and best-fitting parameters.

**Figure 6.**
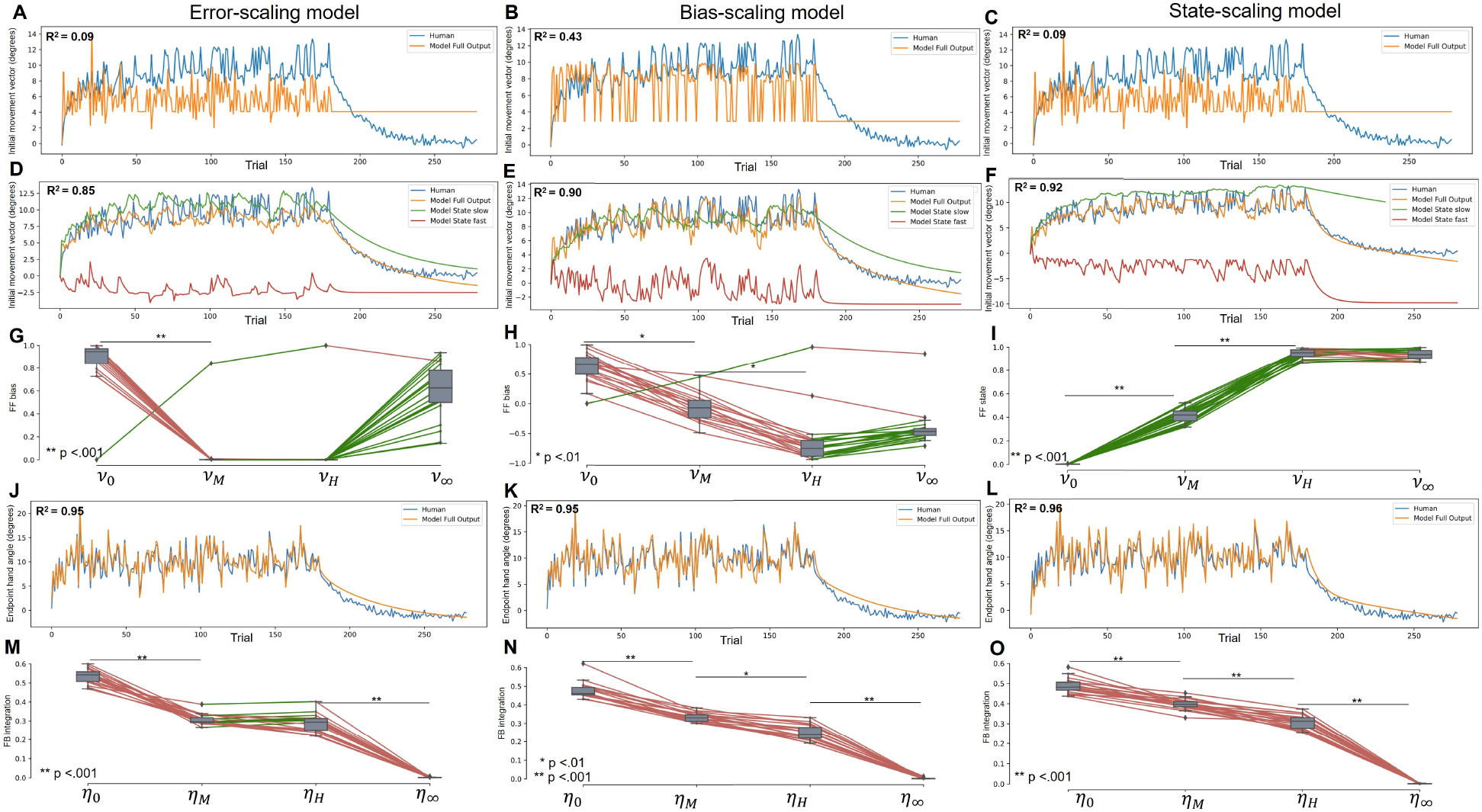
Panels A-C show initial movement vectors averaged across participants in Experiment 2 overlaid with that predicted by the (A) one-state error-scaling model, (B) one-state bias-scaling model, and (C) one-state state-scaling model. Here, human performance averaged across participants is shown in blue and model predictions are shown in orange. Panels D-F show initial movement vectors averaged across participants overlaid with that predicted by the (D) two-state error-scaling model, (E) two-state bias-scaling model, and (F) two-state state-scaling model. Here green and red traces show averaged slow and fast model components separately, which combine to produce the full model predictions in orange. Panels G-I show the distribution of best fitting **ν** parameters for the (G) two-state error-scaling, (H) two-state bias scaling, and (I) two-state state-scaling model. Here, individual differences between parameters are represented by a single colored lines that are green when parameter values increase and red when they decrease. Panels J-L show endpoint hand angles averaged across participants overlaid with that predicted by the (J) two-state error-scaling model, (K) two-state bias-scaling model, and (L) two-state state-scaling model. Here, human performance averaged across participants is shown in blue and model predictions are shown in orange. Panels M-O show the distribution of best fitting **η** parameters for the (M) two-state error-scaling, (N) two-state bias-scaling, (O) two-state state-scaling model.

As seen in Experiment 1, model comparison via Δ*BIC* reveals that the single-state counterparts of each model perform significantly worse than the two-state. Single-state vs two-state error-scaling (Δ*BIC* = −296.84 ± 3.4, *t*(38) = −88.1, *p* > .001, *d* = −27.9). Single-state vs two-state bias-scaling (Δ*BIC* = −239.7 ± 5.5, *t*(38) = −43.4, *p* > .001, *d* = −9.2). Single-state vs two-state state-scaling (Δ*BIC* = −357.9 ± 3.5, *t*(38) = −101.8, *p* > .001, *d* = −32.2).

As in Experiment 1, the slow system state variable (green line in panels A-C) carries the overall adaptation envelope across trials, and the fast system hovers around zero with large fluctuations across trials. Also as seen in Experiment 1, the error-scaling model (*R*^2^ = 0.85) accounts for less of the variance in the behavioural data than either the bias-scaling (*R*^2^ = 0.90) or the state-scaling model (*R*^2^ = 0.92).

Comparing the two-state models Δ*BIC* provides very strong evidence that the error-independant models (state-scaling and bias-scaling models together) account for all 20 individual subject data-sets significantly better than the error-scaling model (state-scaling vs error-scaling Δ*BIC* = −50.9 ± 5.0, *t*(38) = −11.5, *p* < .001, *d* = −3.1; bias-scaling vs error-scaling Δ*BIC* = −41.8 ± 5.4, *t*(38) = −9.4, *p* > .001, *d* = −2.4). When comparing the state-scaling and bias-scaling models, the state-scaling model accounts marginally better for the data in 16 subjects, while the bias-scaling model accounts marginally better for the data in 4 subjects (state-scaling vs bias-scaling Δ*BIC* = −9.1 ± 4.8, *t*(38) = −1.8, *p* = .433, *d* = −0.59).

A one-way ANOVA with best-fitting *ν* values from the state-scaling model as the factor indicates that sensory uncertainty influenced the state term in the feedforward update (*F* = 2765.69, *p* < .001, *ω*^2^ = 0.98). Posthoc t-tests showed that sensory uncertainty inversely scaled the feedforward adaptive update (*ν*_0_-*ν_M_* : *t*(38) = 10.9, *p* < .001, *d* = 2.4; *ν_M_-ν_H_* : *t*(38) = 21.1, *p* < .001, *d* = 4.7). As seen in Experiment 1, the no-feedback *ν_∞_* parameter is not significantly different to the high-uncertainty condition (*ν_H_-ν_∞_* : *t*(38) = 0.96, *p* = 0.22, *d* = 1.46). As observed in Experiment 1, this pattern across feedforward *ν* parameters is also present in the error-scaling and bias-scaling models.

In summary, the state-scaling and the bias-scaling models provide a significantly better fit to the behavioral data than the error-scaling model. The addition of matched endpoint feedback did not change the general finding. The presence of feedback integration fundamentally alters how feedforward adaptation is affected by sensory uncertainty and rules out the possibility that feedforward adaptation is influenced by sensory uncertainty differently depending on the temporal and / or spatial proximity of the sensory feedback signal to movement offset.

Finally, following the results of Experiment 1, feedback integration scales inversely with sensory uncertainty. In particular, the feedback integration aspect of the model lead to *R*^2^ = 0.96 in model endpoint compared to human endpoint data. A one-way ANOVA with best-fitting *η* values (from the bias-scaling model) as the factor confirms that there are differences in how much feedback was integrated across different uncertainty levels (*F* = 1441.1, *p* < .001, *ω* = 0.94). Posthoc t-tests indicate that indeed *η*_0_ > *η_M_* > *η_H_* > *η_∞_* (*η*_0_ – *η_M_* : *t*(38) = 14.9, *p* < .001, *d* = 4.7; *η_M_* – *η_H_* : *t*(38) = 14.9, *p* < .001, *d* = 1.58; *η_H_* – *η_∞_* : *t*(38) = 39.1, *p* < .001, *d* = 8.8).

### Experiment 3

The combined results of Experiment 1 and Experiment 2 show that when feedback integration occurs during a movement, feedforward adaptation depends on sensory uncertainty in a fundamentally different way than when feedback integration is not present. Critically, the error-scaling model – which captures the core assumptions made by all current models of how sensory uncertainty influences feedforward adaptation – does not provide the best account of Experiment 1 and 2. Rather models that assume that sensory uncertainty acts on non-error terms (e.g., bias-scaling and state-scaling) appear to offer a better account of our data.

In Experiment 3, we sought to further probe how sensory uncertainty influences feedback and feedforward error-correction systems. In particular, Experiment 3 disassociates the sensory uncertainty experienced at midpoint from that experienced at endpoint (see Fig. 2C). This allows us to investigate if sensory uncertainty at midpoint dominates sensory uncertainty at endpoint (as might be expected due to the feedback correction made at that time point), or if endpoint dominates midpoint (as might be expected due to the temporal proximity to movement offset) [23, 25].

Figure 7 shows group-averaged initial movement vectors per trial color coded such that the color of the point at trial *t* indicates the sensory uncertainty trial type on trial *t* – 1. There is a clear stratification between conditions according to endpoint uncertainty, regardless of midpoint uncertainty. Furthermore, similar to Experiments 1 and 2, we see that in the transition from lower endpoint uncertainty trials to higher endpoint uncertainty trials (e.g., *σ*_0_, *σ*_0_ → *σ*_0_, *σ_H_*; *σ_H_*, *σ*_0_ → *σ_H_, σ_H_*) the change in movement vector is anti-adaptive.

**Figure 7.**
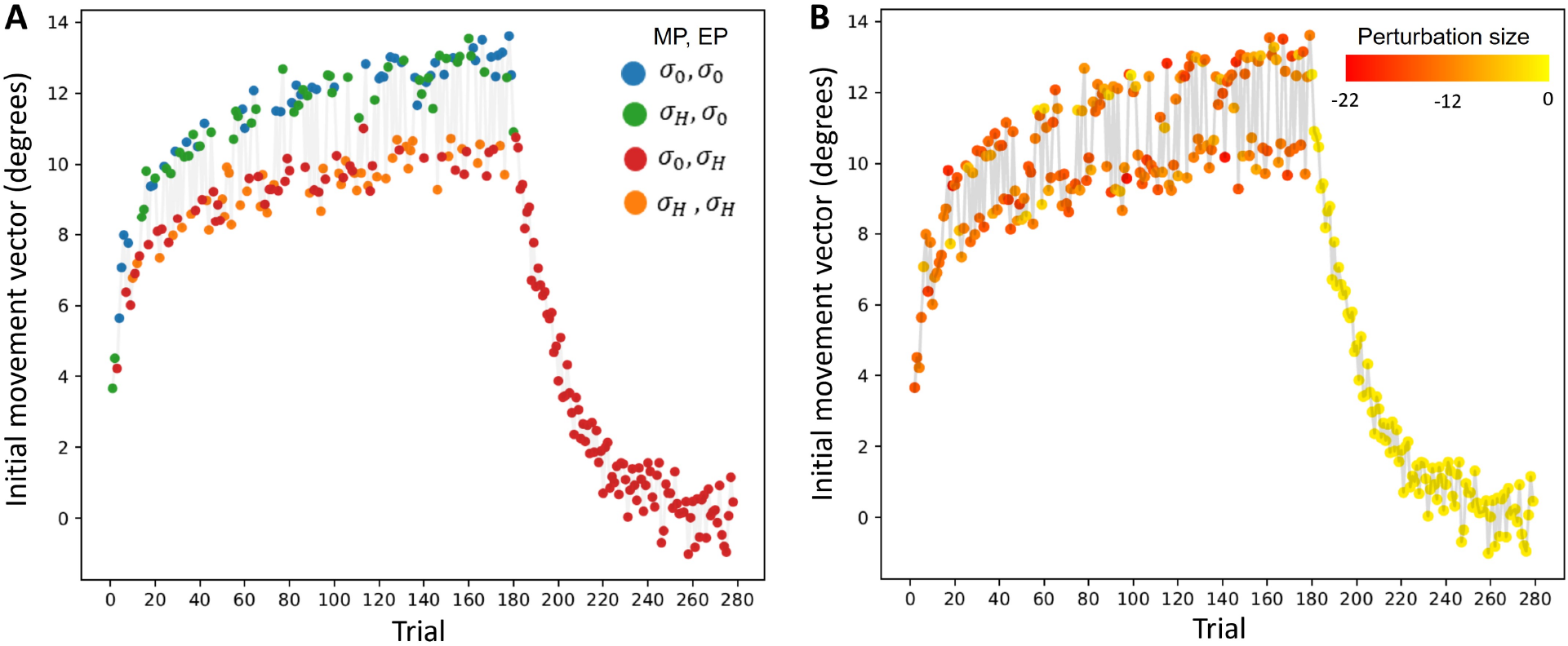
Mean initial movement vector across participants per trial in Experiment 3. (A) The color of the marker on trial t represents one of four midpoint/endpoint combinations of sensory uncertainty applied on the previous trial t – 1 (σ_0_, σ_0_ in blue, σ_H_, σ_0_ in green, σ_0_, σ_H_ in red and σ_H_, σ_H_ in orange). (B) The color of the marker on trial t represents the perturbation applied on the previous trial t – 1.

Figure 8 shows a comparison of one-state and two-state model variants. Here, the behavioral data is overlaid with model predictions and best-fitting feedforward and feedback parameters. Panals A-C show initial movement vectors overlaid with one-state model predictions. All one-state models fit poorly to the behavioural data as seen in experiments 1 and 2. Panels D-R shows a comparison of two-state model variants. Here, the behavioral data overlaid with model predictions and best-fitting parameters. The general results are analogous to those from Experiments 1 and 2. As in Experiments 1 and 2, the slow system state variable (green line in panels D-F) carries the overall adaptation envelope across trials, and the fast system hovers around zero with large fluctuations across trials. The error-scaling model (*R*^2^ = 0.89) accounts for less variance in the behavioural data than either the state-scaling and bias-scaling models model (*R*^2^ = 0.95 for each).

**Figure 8.**
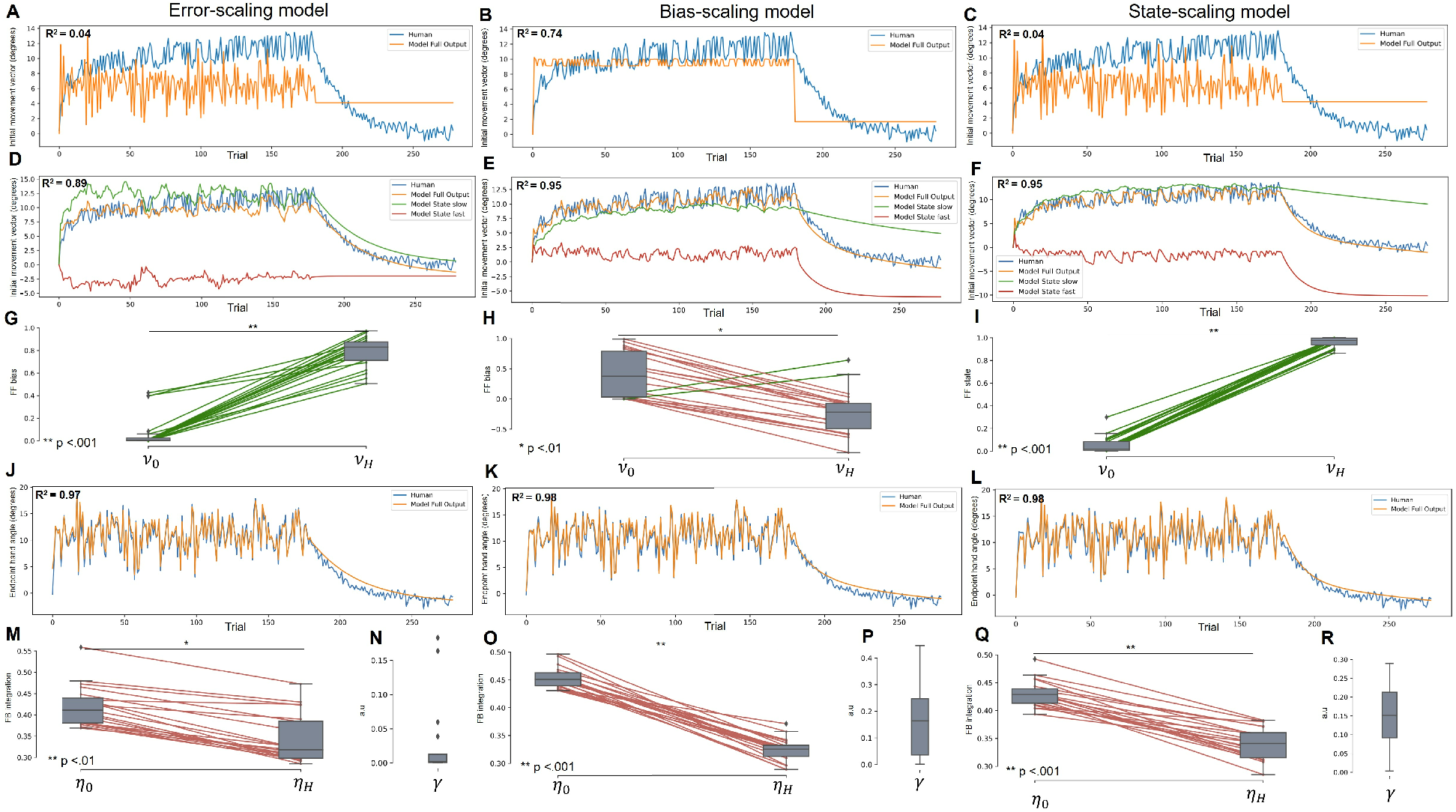
Panels A-C show initial movement vectors averaged across participants in Experiment 3 overlaid with that predicted by the (A) one-state error-scaling model, (B) one-state bias-scaling model, and (C) one-state state-scaling model. Here, human performance averaged across participants is shown in blue and model predictions are shown in orange. Panels D-F show initial movement vectors averaged across participants overlaid with that predicted by the (D) two-state error-scaling model, (E) two-state bias-scaling model, and (F) two-state state-scaling model. Here green and red traces show averaged slow and fast model components separately, which combine to produce the full model predictions in orange. Panels G-I show the distribution of best fitting **ν** parameters for the (G) two-state error-scaling, (H) two-state bias scaling, and (I) two-state state-scaling model. Here, individual differences between parameters are represented by a single colored lines that are green when parameter values increase and red when they decrease. Panels J-L show endpoint hand angles averaged across participants overlaid with that predicted by the (J) two-state error-scaling model, (K) two-state bias-scaling model, and (L) state-scaling model. Here, human performance averaged across participants is shown in blue and model predictions are shown in orange. Panels M, O, and Q show the distribution of best fitting **η** parameters and panels N, P, and R show the temporal discount parameter **γ** for the two-state error-scaling, two-state bias-scaling and two-state state-scaling model, respectively.

Model comparison via Δ*BIC* reveals that the single-state counterparts of each model perform significantly worst than the two-state. Single-state vs two-state error-scaling (Δ*BIC* = −321.6 ± 4.6, *t*(38) = −72.3, *p* > .001, *d* = −22.3). Single-state vs two-state bias-scaling (Δ*BIC* = −144.7 ±5.0, *t*(38) = −32.5, *p* > .001, *d* = −9.1). Single-state vs two-state state-scaling (Δ*BIC* = −381.9 ± 3.7, *t*(38) = −104.1, *p* > .001, *d* = −32.9).

Two-state model comparison via Δ*BIC* provides very strong evidence that the state-scaling model best accounts for the data in all 20 participants (state-scaling vs error-scaling Δ*BIC* = −50.2 ± 4.5, *t*(38) = −11.2, *p* > .001, *d* = −3.52; state-scaling vs bias-scaling Δ*BIC* = −50.2 ± 4.4, *t*(38) = −9.1, *p* > .001, *d* = −3.5). Analysis of best fitting feedforward (*ν*) parameters indicate that sensory uncertainty inversely scaled feedforward adaptive update (*ν*_0_-*ν_H_*) : *t*(38) = −52.6, *p* < .001, *d* = −11.7. Further, in Experiment 3, the temporal discount parameter (*γ*) provides a means by which to determine which of the competing feedback signals (midpoint vs endpoint) principally drives the feedforward update. The mean temporal discount parameter for the state-scaling model (Figure 8R) strongly indicates that endpoint feedback is the dominant feedback signal that drives the feedforward update (*γ* = 0.153 ±0.09). The Shapiro-Wilk test did not show a significance departure from normality (*w*(20) = .948, *p* = .339). The temporal discount parameter for the bias-scaling model (Figure 8P) is statistically similar (Δ*γ* = −0.014 ± 0.031, *t*(38) = −0.462, *p* = .857, *d* = −0.126).

As in Experiments 1 and 2, the standard inverse scaling model of feedback integration provided a very good fit to the data (*R*^2^ = 0.98). The best fitting low uncertainty parameter (*η*_0_) is significantly larger than the high uncertainty parameter (*η_H_*) (*t*(38) = 13.1, *p* < .001, *d* = 2.9) which indicate that feedback integration scales with uncertainty at midpoint (Figure 8Q).

## Discussion

The current study is the first to examine how sensory uncertainty influences feedforward adaptation and feedback integration when they co-occur. We find that the response of feedback integration to sensory uncertainty is unaltered by the presence of feedforward adaptation (i.e., the extent to which sensory feedback is integrated into an ongoing reach is inversely scaled by its level of uncertainty regardless of the presence or absence of feedforward adaptation [14, 15, 16, 26]). However, in sharp contrast to studies examining these processes in isolation of each other – all of which have found that sensory uncertainty inversely scales an error-driven update [14, 15, 17, 18, 19, 20, 21, 27] – we show that sensory uncertainty punctuates a slow envelope of error reduction with large and abrupt changes to initial movement vectors that are insensitive to the magnitude and direction of the sensed movement error. A key difference between these earlier studies and the present one is that these earlier studies examined feedback integration and feedforward adaptation in isolation from one another, whereas our study examines them as they co-occur.

### Divergence from existing work

Standard models of motor learning assume a linear relation between adaptation rate and error size [9, 27, 28, 29, 30, 31, 32], and the influence of sensory uncertainty on this process has been thought to inversely scale the error-driven update [18, 19, 22, 33, 34] (i.e., our error-scaling model). These results have often been interpreted through the lens of Bayesian [33] or other optimality frameworks such as that of Kalman filters [35]. These frameworks assume that as sensory uncertainty increases the motor system should limit adaptation in response to observed errors because they likely reflect higher sensory noise (to which adaptation would be sub-optimal) instead of actual changes in the external environment (to which adaptation would be optimal). Consequently, when sensory uncertainty is high, the motor system should adapt less to a given error. For instance, compare two high uncertainty conditions, one in which a large perturbation was applied (e.g., 22°), and one in which a small perturbation was applied (e.g., 12°). In each case, the effect of uncertainty should be the same, involving a down-weighting of the sensed error by the same amount. However, the overall amount of adaptation should be different because it is scaled by error size.

The pattern observed in our data does not clearly resonate with the above description. In particular, initial movement vectors following high sensory uncertainty trials were always a few degrees less than they were on the last low sensory uncertainty trial, regardless of the magnitude or even the direction of the just experienced movement error. Motivated by this apparent insensitivity to error, we developed a bias-scaling model in which sensory uncertainty scales a constant bias term in the feedforward update, and a state-scaling model in which sensory uncertainty scales the rate that the system returns to baseline. In both the bias-scaling and the state-scaling models, sensory uncertainty has no effect on the error-driven component of the feedforward update. In every experiment – and even in every participant – the bias-scaling and state-scaling models provided better fits than the error-scaling model. This is a significant point of divergence from the existing literature.

### Adaptation versus aiming

Our results can be viewed from at least two perspectives. From one perspective, all observed initial movement vectors reflect the true adapted state of the motor system. Accordingly, adaptation following low sensory uncertainty trials occurs more or less normally, which is to say in a direction that tends to reduce the error likely to be experienced on future trials. But trials following greater sensory uncertainty exhibit a rather strange pattern – they would tend to increase the error likely to be experienced on future trials – i.e., they would be *anti-adaptive*.

An alternative perspective is that the initial movement vectors following greater uncertainty trials reflect the true adapted state of the motor system, and another factor drives an abrupt change in initial movement vectors following low sensory uncertainty trials. This view is reinforced by the pattern of initial movement vectors observed during the washout phase of all three experiments: initial movement vectors start their decay back to baseline from the current state of the greater uncertainty trials, not from the level of the lower uncertainty trials.

One candidate for what might be driving these abrupt changes is the use of explicit aiming strategies [36, 37, 38, 39]. In essence, low sensory uncertainty trials could provide a clear signal for participants to notice the mismatch between where they were aiming and where the cursor landed, which in turn could be used to form a hypothesis about the perturbation magnitude and direction. Following greater sensory uncertainty trials, confidence in this hypothesis might be eroded by the poor sensory feedback, and participants might simply reach straight to the target (or as straight as their current level of adaptation allows) in an attempt to get a better explicit estimate of the true perturbation.

This possibility is broadly consistent with the results of Experiment 3, which clearly show that feedforward adaptation stratifies according to sensory uncertainty at endpoint regardless of the sensory uncertainty experienced at midpoint. Implicit motor adaptation is thought to be primarily driven by sensory prediction errors [13, 40] (i.e., the difference between the expected and observed sensory consequences of a motor command), which can be provided by both midpoint and endpoint feedback. Strategy learning on the other hand is sensitive to task performance errors [36, 37, 41] (i.e., the difference between the task goal and the observed outcome), which may be preferentially encoded by endpoint feedback. Findings from Experiment show that endpoint feedback, and in turn perhaps explicit aiming strategies keyed to task performance errors, are the driving force behind our data.

We should, however, be careful not to endorse this possibility too quickly. First, the best fitting models in our experiments (i.e., the state-scaling and the bias-scaling models) provided good fits to our data without appealing to aiming strategies or otherwise distinguishing between sensory prediction errors and task performance errors. Second, it is also possible that sensory uncertainty at both midpoint and endpoint jointly influence feedforward adaptation, and it is only the specific conditions used in Experiment 3 that failed to detect an effect of midpoint uncertainty. For instance, the influence of sensory uncertainty on adaptation may reflect both its level of visual blur and its temporal proximity to movement offset [23, 24, 25]. Increased sensitivity to error information near the time of movement offset is also supported by evidence that complex spike discharge in the cerebellum, hypothesized to encode motor error, occurs around 100ms after error feedback is provided [42, 43].

Ultimately, the experiments reported here were not optimally designed to adjudicate between these two possible explanations. To dissociate the contributions of implicit adaptation and explicit strategies on reaching behavior, future studies could employ pre-reach aiming reports [37, 41, 44] or clamped visual feedback that is invariant for all hand movements [37, 40, 41, 44, 45, 46].

### Can paradigm differences explain our divergent results?

There are three features of our paradigm that may explain why our results diverge from the existing literature. First, we used interleaved uncertainty conditions whereas most other relevant studies use blocked designs [15, 21, 22, 47]. However, it is unlikely that our interleaved design accounts for the shift from an error-dependent to an error-independent effect, since Wei and Körding (2010) applied randomly interleaved levels of sensory uncertainty across trials and an error-scaling model was well suited to capture their results [19].

Second, we used a stochastic perturbation that varied from trial-to-trial, whereas most other studies use a fixed perturbation. To our knowledge, Körding and Wolpert [14] is the only other study that varied both the sensory uncertainty and the imposed perturbation on a trial-by-trial basis. Although they report on feedback integration of sensory uncertainty after thousands of trials of adaptation, they do not report initial movement vectors from the beginning of the experiment where little to no adaptation would have occurred. Because of this we cannot directly compare our feedforward adaptation results with their findings.

The remaining possibility is that our divergent results reflect the fact that all previous studies intentionally isolate feedforward and feedback processes by presenting endpoint feedback only [17, 18, 19, 20, 21, 22]. In these designs, endpoint feedback is safely assumed to drive feedforward adaptation in the absence of feedback corrections. By contrast, our experiments were explicitly designed to investigate the influence of sensory uncertainty when feedback integration and feedforward adaptation co-occur. Future research should seek to identify how the interaction of these two processes leads to qualitatively different behavior when they are studied alone than when they are studied together.

## Materials and Methods

### Participants

A total of 60 naive subjects (32 males, 28 females, age 17-33 years) with normal or corrected to normal vision and no history of motor impairments participated in the experimental study. All subjects gave informed consent before the experiment and were either paid and recruited from the [redacted] Cognitive Science Participant Register or were [redacted] undergraduates participating for course credit. All experimental protocols were approved by the [redacted] Human Research Ethics Committee (protocol number: 52020339922086). Subjects were randomly assigned to one of three experiments (n=20 per experiment). Sample sizes were consistent with field-standard conventions for visuomotor adaptation experiments [48, 49, 50].

### Experimental Apparatus

A unimanual KINARM endpoint robot (BKIN Technologies, Kingston, Ontario, Canada) was utilized in the experiments for motion tracking and stimulus presentation (figure 1). The KINARM has a single graspable manipulandum that permits unrestricted 2D arm movement in the horizontal plane. A projection-mirror system enables presentation of visual stimuli that appear in this same plane. Subjects received visual feedback about their hand position via a cursor (solid white circle, 2.5 mm diameter) controlled in real-time by moving the manipulandum. Mirror placement and an opaque apron attached just below the subject’s chin ensured that visual feedback from the real hand was not available for the duration of the experiment.

### General Experimental Procedure

Subjects were instructed to perform fast and accurate horizontal reaches using their dominant hand. In our sample, this was the right hand for all participants. Subjects were also instructed to use cursor feedback, whenever it was available, to guide their movement.

Subjects performed reaches from a starting position located at the center of the workspace (solid green circle, 0.5cm in diameter) to a single reach target (solid red circle, 0.5 cm in diameter) located 10 cm away. When subjects moved the cursor within the boundary of the start target its color changed from red to green and the reach target appeared, indicating the start of a trial. Subjects were free to reach at any time after the start target color changed. Subjects first completed a 20 trial baseline phase during which veridical online feedback was provided. Immediately following baseline, a 180 trial adaptation phase was completed. During the adaptation phase, once the cursor exited the start target, cursor feedback was extinguished and rotated counterclockwise (to the left) of the true hand position by an amount drawn at random on each trial from a Gaussian distribution with a fixed mean of 12° and standard deviation of 4°. Random trial-by-trial perturbations, the order of which was trial-matched across all subjects, was applied in order to avoid a predictable feedback command during the adaptation phase and to provide a means by which to capture the effect of sensory uncertainty at a trial-by-trial resolution.

Contingent upon the specific experiment (see descriptions below for details), displaced cursor feedback was provided at reach midpoint (100ms duration) and/or at endpoint (100ms duration) or withheld altogether. To help guide the participant’s hand back to the starting position a green ring centered over the starting position appeared with a radius equal to the distance between the hand and starting position. Once the participant’s hand was 1 cm from the starting position the ring was removed and cursor feedback was reinstated.

To investigate the effect of sensory uncertainty on feedback integration and feedforward adaptation, the uncertainty of the visual information provided about the visuomotor perturbation (true cursor position) was varied as described below. One of four visual uncertainty levels (*σ*_0_, *σ_M_*, *σ_H_*, *σ*_∞_) were selected and applied on a given trial according to the specific experimental protocol, with trial sequence matched across subjects. In the zero uncertainty condition (*σ*_0_), feedback was a single white sphere (0.5 cm in diameter), identical to the initial cursor. In the moderate uncertainty condition (*σ_M_*), feedback was one of ten randomly generated point clouds comprised of 50 small translucent white spheres (0.1 cm in diameter) distributed as a two-dimensional Gaussian with a SD of 0.5cm and a mean centered over the true (displaced) cursor position on the current trial. In the high uncertainty condition (*σ_H_*), everything was the same as the moderate uncertainty condition (*σ_M_*) except that the point clouds had a SD of 1 cm. In the unlimited uncertainty condition (*σ*_∞_), no feedback was provided at all.

As a means by which to investigate adaptation aftereffects, immediately proceeding the adaptation phase, a 100 trial washout phase was completed during which no cursor feedback was provided during the reach. Subjects were instructed to move their hand straight through the target as accurately as possible.

The maximum allowable time to complete a reach was 1000 ms. Irrespective of the cursor’s position, if subjects did not cross the lower bound of the end target radius (9.5cm) the trial would time out and restart.

### Experiment 1

All four feedback uncertainty types (*σ*_0_, *σ_M_*, *σ_H_*, *σ_∞_*) were applied on 25% of trials (45 trials each) at midpoint only (figure 2a). No feedback was provided at endpoint. The order of uncertainty conditions and perturbation values were randomised and trial-matched across all subjects.

### Experiment 2

The protocol employed in Experiment 2 was identical to Experiment 1 except for one key difference: both midpoint and endpoint feedback – matched in uncertainty level – were provided on each trial (figure 2b).

### Experiment 3

Experiment 3 consisted of four trial types (figure 2c). Trial type 1 consisted of low uncertainty midpoint and low endpoint feedback (*σ*_0_*, σ*_0_), trial type 2 consisted of low uncertainty midpoint and high uncertainty endpoint feedback (*σ*_0_, *σ_H_*), trial type 3 consisted of high uncertainty midpoint and low uncertainty endpoint feedback (*σ_H_, σ*_0_), and trial type 4 consisted of high undertainty midpoint and high uncertainty endpoint feedback (*σ_H_, σ_H_*). Each of the four trial types occurred on 25% of trials (45 trials each). The perturbation applied to each trial was randomly drawn from a normal gaussian distribution with a mean of 12°and standard deviation of 4°counter clockwise. The trial order of uncertainty conditions and perturbation values were randomised and matched across subjects.

### Data Analysis

Movement kinematics including hand position and velocity were recorded for all trials using BKIN’s Dexterit-E experimental control and data acquisition software (BKIN Technologies). Data was recorded at 200 Hz and logged in Dexterit-E. Custom scripts for data processing were written in MATLAB (R2013a). Data analysis and model fitting was done in Python (3.7.3) using the numpy (1.19.2) [51], SciPy (1.4.1) [52], pandas (1.1.3) [53], matplotlib (3.3.2) [54], and pingouin (0.3.11) [55] libraries. Best fitting model parameter distributions were analysed using analysis of variance (ANOVA) and Welch t-tests (*α* < .05). We report Δ*BIC* and compare the *BIC* distributions for model comparison via Dunnett’s post-hoc test and correct for multiple comparisons using the Bonferroni correction.

A combined spatial- and velocity-based criterion was used to determine movement onset, movement offset, and corresponding reach endpoints [56, 57]. Movement onset was defined as the first point in time at which the movement exceeded 5% of peak velocity after leaving the starting position. Movement offset was similarly defined as the first point in time at which the movement dropped below 5% of peak velocity after a minimum reach of 9.5 cm from the starting position in any radial direction, and reach endpoint was defined as the (*x, y*) coordinate at movement offset. During the no-feedback washout phase, endpoint hand-angle was analysed to investigate adaptation aftereffects.

### Computational modelling

Here, we develop three models of how sensory uncertainty influences feedforward adaptation and feedback integration. At a high level, each model is characterised by the following attributes:

- A feedforward motor plan is computed at movement onset that is an attempt to reach in a straight line from the starting position to the target location, and a feedback motor command is computed at movement midpoint that is an attempt to correct the ongoing movement for any error experienced at midpoint.
- Feedforward motor plans are adapted on a trial-by-trial basis using both the error experienced at midpoint and the error experienced at endpoint as a learning signal.
- The gain applied to feedback corrections is similarly adjusted on a trial-by-trial basis but is sensitive only to the error experienced at endpoint.
- The sensory uncertainty experienced at midpoint and / or endpoint modulates the between-trial feedforward update, and the within-trial feedback correction, but – for simplicity – not the between-trial feedback gain update.

The feedforward adaptation component of all three models is based on simple discrete-time linear dynamical systems – so-called state-space models [30]. The simplest version of these models assumes that an internal state variable *x* maps desired motor goals to motor plans *y*, and that *x* is updated on a trial-by-trial basis in response to sensory feedback about movement error. The update to *x* has (1) an error term that determines how the internal state is updated after a movement error is detected, (2) a bias term that determines the baseline mapping that will be returned to in the absence of sensory input and (3) retention term that determines how quickly the internal state returns to baseline after sensory feedback about error is removed. This arrangement is encapsulated in the following equations:

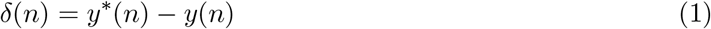

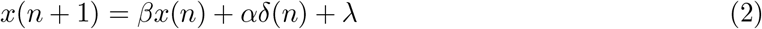

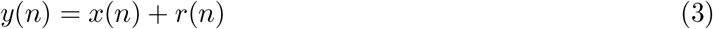

where *n* is the current trial, *δ*(*n*) is the error (i.e., the angular distance between the reach endpoint and the target location), *y*\*(*n*) is the desired output (e.g., the angular position of the reach target), *y*(*n*) is the motor output and corresponds to the angle of the movement that will be generated when trying to reach to the target (i.e., it is a readout of the sensorimotor state), *x*(*n*) is the state of the system (i.e., the sensorimotor transformation), *β* is a retention rate that describes how much is retained from the value of the state at the previous trial, *α* is a learning rate that describes how quickly states are updated in response to errors, *λ* is a constant bias, and *r*(*n*) is the imposed rotation.

These models are sometimes equipped with a second internal state variable [36, 45] as follows:

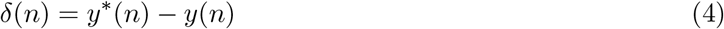

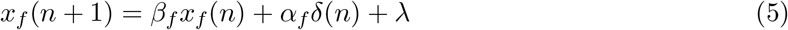

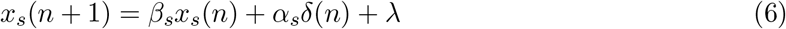

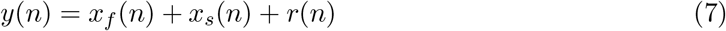

where *x_f_* is a *fast* state variable, *x_s_* is a *slow* state variable, *β_f_* < *β_s_*, and *α_f_* > *α_s_*. That is, feedforward adaptation is assumed to arise from the combination of a slow-but-stable system and a fast-but-labile system. Existing studies of have not strongly delineated the appropriateness of one-state versus two-state models in mapping out how sensory uncertainty influences feedforward adaptation, and as such, we explore both one-state and two-state model variants in this paper.

The simple state-space framework just described assumes motor output reflects the execution of feedforward motor commands, whereas the behavior observed in our experiments is also likely influenced by feedback motor commands. We therefore augment the simple models presented above as follows.

The total motor output of the model is defined at three discrete time points within each trial. We denote the time of reach initiation as *t*_0_, the time of midpoint crossing as *t*_MP_, and the time of endpoint crossing as *t*_EP_. The total motor output on trial *n* at any time *t* denoted *y*(*n, t*) is a combination of feedforward *y*_ff_(*n*) and feedback *y*_fb_(*n, t*) motor commands as follows:

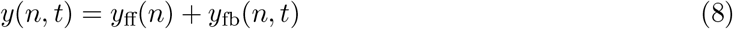

Note that feedforward motor output is not a function of time within a trial because we assume that the feedforward motor output is computed at *t*_0_ and remains fixed throughout the rest of each trial. This is equivalent to assuming that the execution of the movement occurs too rapidly for new feedforward motor planning to influence the ongoing movement.

In the *single-state* models, the feedforward motor command *y*_ff_(*n*) is determined by a single internal state variables denoted by *x*_ff_(*n*) that maps the current movement goal to motor commands as follows:

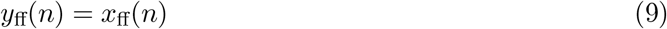

In the *two-state* models, the feedforward motor command *y*_ff_(*n*) is determined by two internal state variables denoted by *x*_ff_*f*__(*n*) and *x*_ff_*s*__(*n*) that map the current movement goal to motor commands as follows:

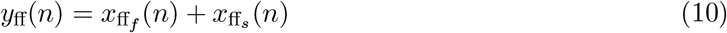

At reach initiation, sensory feedback has not yet been provided so the feedback motor command is zero:

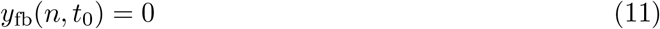

If sensory feedback is provided at midpoint, then the following sensory prediction error is experienced:

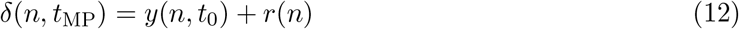

Here, *δ*(*n, t*_MP_) is the sensory prediction error, and *r*(*n*) is the visuomotor rotation applied on trial *n*. Notice that the motor command issued at time *t*_0_ is responsible for generating the sensory prediction error at time *t*_MP_. In response to this sensory prediction error, the following compensatory feedback motor command is triggered:

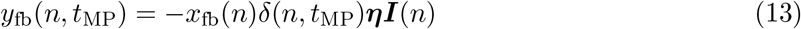

Here, *x*_fb_(*n*) is an internal state variable that represents the gain of the feedback controller, ***η*** = [*η*_0_, *η_M_*, *η_H_*, *η*_∞_] is a row vector of free parameters encoding the sensory uncertainty of the midpoint feedback (one value for each possible level of sensory uncertainty), and ***I***(*n*) is a column vector that indicates what level of midpoint uncertainty was present on trial *n*. Notice that the feedback motor command is just some proportion of the experienced error in magnitude and in the opposite direction – because of the leading negative sign – hence it serves to reduce movement error.

If endpoint sensory feedback is provided, the following sensory prediction error is experienced:

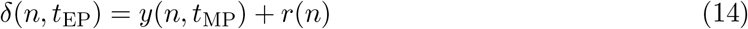

Notice that the motor command issued at time *t*_MP_ is responsible for generating the sensory prediction error at time *t*_EP_. In the transition from trial *n* to trial *n* + 1, the gain of the feedback controller is updated in response to this sensory prediction error as follows:

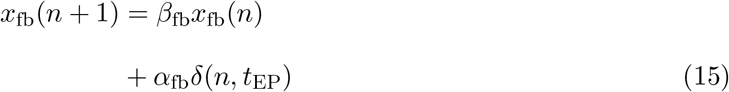

Note that we assume that updates to feedback gain are not sensitive to sensory uncertainty.

We built several models each making different assumptions about how updates to the feedforward state(s) are determined. In particular, we built (1) *error-scaling* models which assume that sensory uncertainty scales the contribution of the error term (e.g., the *αδ(n*) in eq. 3), (2) *state-scaling* models which assume that sensory uncertainty scales the contribution of the retention term (e.g.,the *βx*(*n*) in eq. 3), and (3) *bias-scaling* models which assume that sensory uncertainty scales the contribution of the bias term (e.g.,the *λ* in eq. 3).

In the *two-state* version of these models, we assume that sensory uncertainty only influences *x*_ff_*f*__. That is, we assume that *x*_ff_*s*__ is independent of sensory uncertainty. In particular, the feedforward internal state *x*_ff_*s*__ is updated between trials in response to the sensory prediction errors experienced at midpoint and at endpoint as follows:

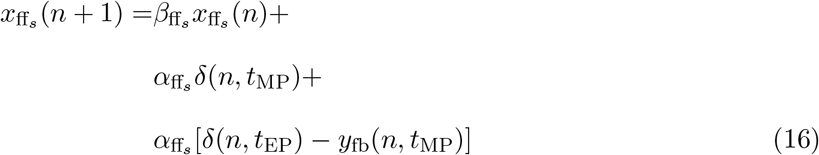

Here, *α*_ff_*s*__ is a learning rate parameter and *β*_ff_*s*__ is a retention parameter, both bound between [0, 1]. Notice that the feedback command issued at midpoint is taken to be an error signal in these equations – in the term [*δ*(*n, t*_EP_) – *y*_fb_(*n, t*_MP_)] – which is a common assumption in models that join feedforward and feedback control [58, 59, 60, 61].

#### Error-scaling models

The uncertainty of sensory feedback influences the update to *x*_ff_ in the *single-state error-scaling* model by acting as a gain on the learning rate *α*_ff_. The update is given by:

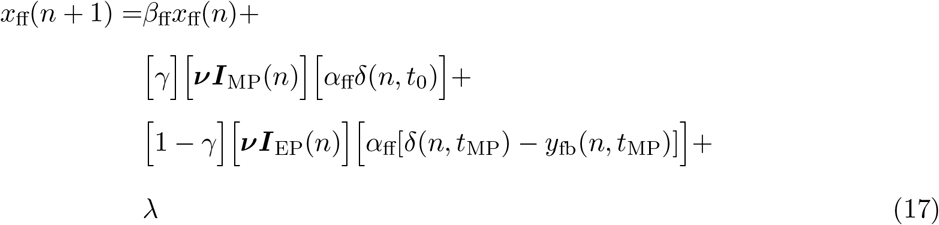

Here, ***ν*(*n*)** = [*ν*_0_, *ν_M_*, *ν_H_*, *ν*_∞_] is a row vector of free parameters (one value for each level of sensory uncertainty), with boundary conditions between [0, 1], to represent the scaling effect of sensory uncertainty. ***I***_MP_(*n*) is a column vector that indicates the uncertainty of sensory feedback that was present on trial *n* at midpoint, ***I***_EP_(*n*) is a column vector that indicates the uncertainty at endpoint, and *γ* is a temporal discounting parameter, bound between [0, 1], that determines the relative weighting of midpoint versus endpoint feedback on the overall state update. For instance, if *γ* > 0.5, midpoint feedback drives the majority of the state update. If *γ* < 0.5, endpoint feedback drives the majority of the state update. Note that any interpretation of the temporal discount parameter is relevant only in the case of Experiment 3, which is the only paradigm that applies unmatched midpoint and endpoint feedback, a feature required to demarcate the effect of the learning rate parameter from the effect of the temporal discounting parameter. The constant bias term *λ* is bound between [−10, 10]. The update to *x*_ff_*f*__ in the *two-state error-scaling* follows exactly the same equation, while the update to *x*_ff_*s*__ is denoted by equation 16.

The classic finding that sensory uncertainty inversely scales the magnitude of the error-driven component of the feedforward update [15, 17, 18, 19, 20, 21, 22] would be recapitulated here if (1) the error-scaling model provides the best-fitting account of our data, and (2) the best-fitting parameters were such that *ν*_0_ > *ν_M_* > *ν_H_* > *ν*_∞_.

#### State-scaling models

The uncertainty of sensory feedback influences the update to *x*_ff_ in the *single-state bias-scaling* model by acting as a gain on the retention term *β*_ff_. The update is given by the following equations:

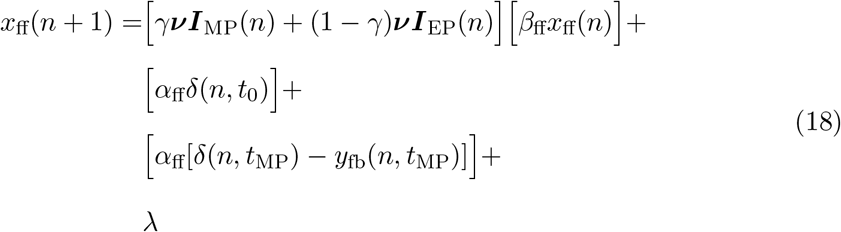

All parameters, nomenclature and bounds are identical to that described above for the error-scaling model. The update to *x*_ff_*f*__ in the *two-state state-scaling* model follows exactly the same equation, while the update to *x*_ff_*s*__ is denoted by equation 16.

#### Bias-scaling models

The uncertainty of sensory feedback influences the update to *x*_ff_in the *single-state state-scaling* model by acting as a gain on the bias term *λ*. The update is given by:

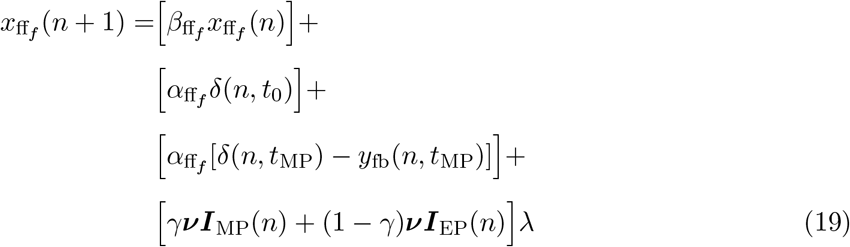

All parameters and nomenclature are identical to that described above for the error-scaling model. The update to *x*_ff_*f*__ in the *two-state bias-scaling* follows exactly the same equation, while the update to *x*_ff_*s*__ is denoted by equation 16.

#### Parameter estimation

For each model, we obtained best-fitting parameter estimates on a per subject basis by minimising the following sum of squared error difference between the observed and predicted midpoint and endpoint hand angles:

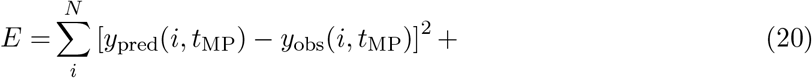

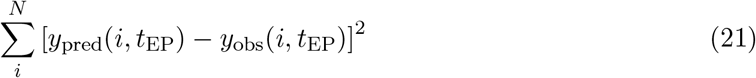

Here, *N* is the number of trials, *y_pred_*(*i,t*_MP_) and *y*_pred_(*i, t*_EP_) are the model predicted hand angle at midpoint and endpoint, respectively, on trial *i*, and *y*_obs_(*i,t*_MP_) and *y*_obs_(*i,t*_EP_) are the corresponding hand angles observed in a human participants. To find the parameter values that minimised *E*, we used the differential evolution optimization [62] method implemented in *SciPy* [52].

For each model, we computed the Bayesian Information Criterion (*BIC*) as follows:

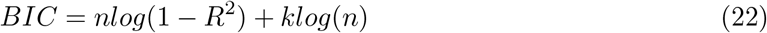

Here *k* represents the number of model parameters for each model, *n* represents the number of data points, and *R*^2^ is the proportion of variance explained by the optimised model. We selected the model with the lowest *BIC* value as best-fitting. We characterised the strength of the evidence for the model with the lowest *BIC* value by computing the difference in *BIC* (Δ*BIC*) between each pair of models on a per subject basis. |Δ*BIC*| < 2 is not significant, 2 < |Δ*BIC*| < 6 is weak, 6 < |Δ*BIC*| < 10 is strong, and |Δ*BIC*| > 10 is very strong [63].

## Code Accessibility

Data and analysis code can be accessed at: https://github.com/hewitsonchris/sensory_uncertainty_2022.git

